# qPaLM: quantifying occult microarchitectural relationships in histopathological landscapes

**DOI:** 10.1101/828004

**Authors:** Timothy J Kendall, Catherine M Duff, Andrew M Thomson, John P Iredale

## Abstract

Optimal tissue imaging methods should be easy to apply, not require use-specific algorithmic training, and should leverage feature relationships central to subjective gold-standard assessment. We reinterpret histological images as landscapes to describe quantitative pathological landscape metrics (qPaLM), a generalisable framework defining topographic relationships in tissue using geoscience approaches. qPaLM requires no user-dependent training to operate on all image datasets in a classifier-agnostic manner to quantify occult abnormalities, derive mechanistic insights, and define a new feature class for machine-learning diagnostic classification.

## Main

Traditional diagnostic pathology remains the gold-standard means of assessing tissue but is a subjective and poorly-reproducible craft. Progress has been made to introduce objectivity and reproducibility into the field by computational interrogation of digital histological images^1^ but there are conceptual short-comings and practical obstacles in current approaches. Current image analysis methods require application-specific algorithm training by end-users, an impediment to widespread use. Further, current approaches are unable to extract understanding from the histological context of observable features, the most critical component informing skilled subjective assessment. We address these short-comings by reinterpreting images as histological landscapes to describe quantitative pathological landscape metrics (qPaLM), a generalisable scale-independent framework that leverages essential feature relationships using geoscience approaches. qPaLM requires no further user-dependent training to operate on image datasets in a classifier-, species-, and disease-agnostic manner within computational workflows. This provides a pathologically intuitive framework that identifies occult abnormalities, derives mechanistic insights, and defines a new feature class for machine-learning disease classification.

A significant impediment to the widespread adoption of current quantitative digital pathology methods is the necessity for study-specific algorithm training. This is time-consuming, and critically precludes inter-study comparison of measured outputs or outcomes in animal modelling of disease or clinical trials. Only by developing methods that can be extensively validated and applied uniformly and intuitively across studies without a need for specialist input can quantitative digital pathology disrupt classical subjective assessment in a research setting or within routine practice.

Feature recognition is central to both traditional and computational methods. Although advances in computational feature annotation using deep-learning methods^2–5^ have increased the accuracy of image segmentation, ‘real-world’ diagnostic acuity is a function of histological literacy -an appreciation of the histological context and relationships between features – rather than accurate feature recognition alone. Understanding from these feature relationships is not currently exploited computationally, representing a significant opportunity for a more creative approach to harness concepts with proven, real-world value. We reasoned that any annotated histological image could be conceptualised as a simple two-dimensional landscape in a generalisable manner to permit quantitation of feature relationships by methods developed for landscape analysis in geosciences and ecology. To this end, we have developed qPaLM, a framework that applies spatial point and categorical landscape pattern analysis in a histopathological context.

The analytic input for qPaLM is a landscape pattern created by manual or computational annotation of a histological image. Complex computational methods to fully classify histological images are available, and their ease-of-use and accuracy continue to increase. The output of classifiers such as U-net^2^ can be a categorical landscape equivalent to those generated in large-scale mapping and geoscience studies. Whilst the scales differ, the fundamental nature of the data representation is the same (Figure 1a). Categorical landscape patterns are mosaics of discrete areas (‘patches’) belonging to defined conditions (‘classes’). In landscape ecology, patches are environmentally homogeneous areas with patch boundaries reflecting the significant change in conditions between them. Conceptually, the histological landscape also consists of a mosaic of ‘environmentally’ similar areas represented by tissues or cells and extracellular microarchitectural structure. Analysis of categorical landscape patterns can generate metrics describing individual patches, the patch class, or defining the landscape as a whole. Class and landscape-level metrics describe the histological topography in a holistic and novel language whilst individual patch-level metrics provide metrics complementary to single-cell/group histological phenotyping^6^ provided by existing methods.

**Figure 1.**
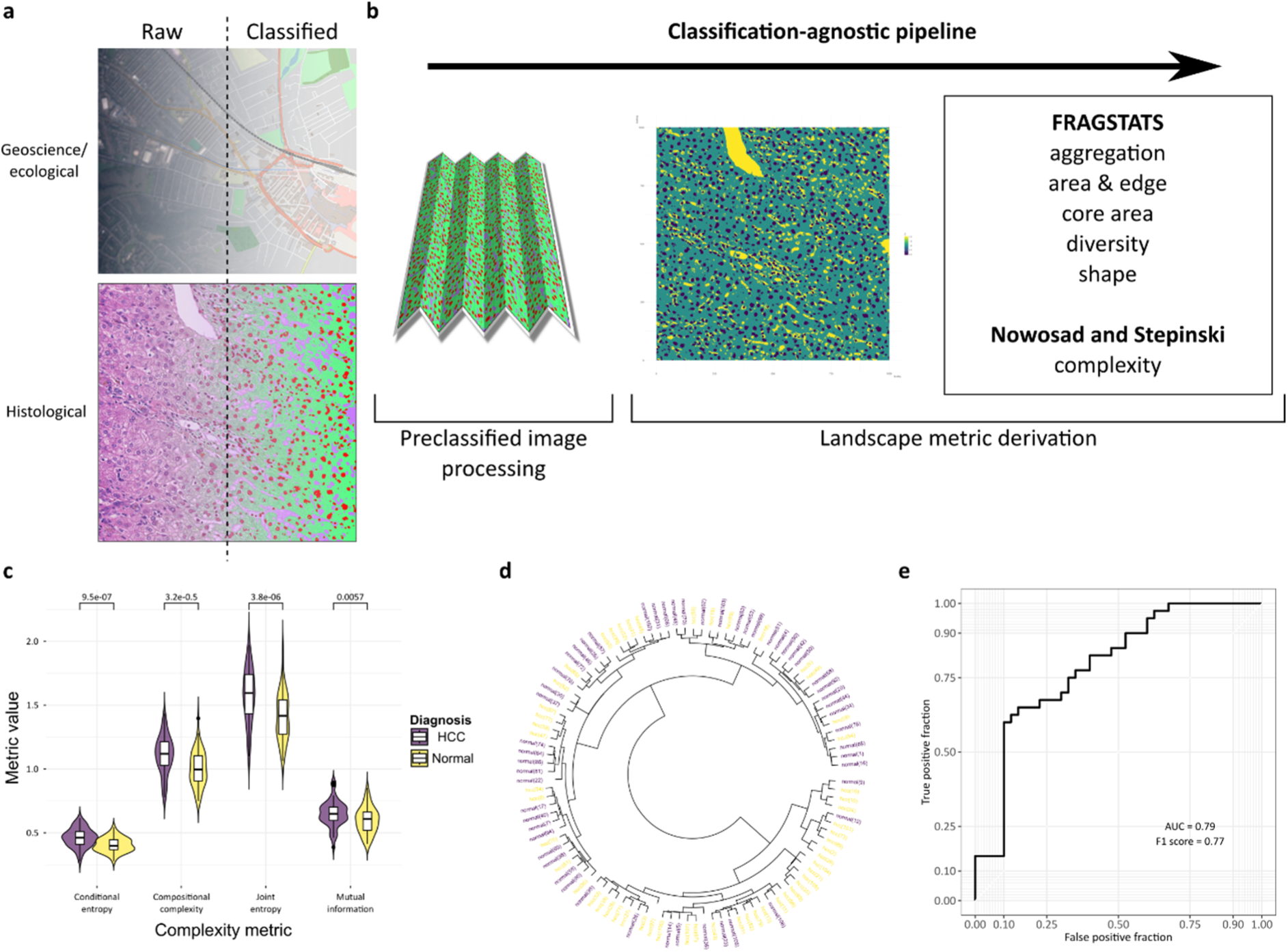
Fully classified histological images can be considered categorical maps and analysed as part of a fully computational pipeline using landscape ecology and geosciences methodologies. A. Categorical representations of the landscape are routinely evaluated in landscape ecology and geosciences and can be evaluated by specific tools. The generation of a fully segmented output image from a histological input, by any available method, is an analogous process differing only in scale. B. A pipeline using fully segmented images generated by a WEKA classifier converts the images to an appropriate file format feeds through the landscapemetrics package in R, generating the complete suite of metrics described in FRAGSTATS, and other holistic landscape measures of complexity and organisation. C. Individual complexity metrics (for example, form lesional liver and hepatocellular carcinoma) can be used as discrete phenotyping measures (violin plot with kernel density and median (centre line), first and third quartiles (lower and upper box limits), 1.5x interquartile range (whiskers); p-values of Welch paired two-sided two-sample t-test for each metric, n=54) or combined to allow unsupervised clustering (D). E. The complete suite of landscape metrics can be used for machine-learning applications such as diagnostic classification (for example, classification as normal or hepatocellular carcinoma, Area Under the Receiver Operating Characteristics curve for diagnostic accuracy on test set).

As proof-of-principle, we trained a basic machine-learning classifier to deconvolve routine haematoxylin and eosin staining in histological images of normal liver and primary liver cancer (hepatocellular carcinoma), creating three classes – nuclei, cytoplasm and vascular channels. We developed a pipeline using the classified images to analyse the landscape patterns using methods derived from the FRAGSTATS suite^7^, a spatial pattern analysis program for categorical maps originally developed in association with the USDA Forest Service, as well as more recently described measures of landscape complexity^8^ (Figure 1b).

The four holistic metrics of landscape complexity^8^ provide single values derived from each landscape. These four metrics alone can be used as quantitative descriptors of the complete histological landscape to describe differences between paired tumour and normal liver, augmenting the subjective diagnosis (Figure 1c). The four metrics in combination were used for unsupervised clustering and provided good disease discrimination (Figure 1d).

The large number of metrics generated from categorical histological landscapes (Supplementary file 1) can be used in downstream applications such as machine-learning driven diagnostic classification. Selected metrics were used as features for model training. A random forest model was constructed from the selected features, and the predictive value of the model determined on a test set (Figure 1e), demonstrating the applicability of this type of landscape feature metrics in pertinent down-stream uses. Examination of variable importance measures in this application indicates that features derived from the ‘nuclear’ class in ‘aggregation’ and ‘area and edge’ categories are the most highly ranked (Supplementary figure 1). These metrics represent nuclear morphology and distribution, critical cytological features used by pathologists to make a subjective diagnosis. Further, the most highly ranked interactions were between nuclear and vascular channel features, indicating that the histological context of nuclei is computationally determined to be critical. This demonstrates that a fully computational landscape approach independently identifies and utilises features used in gold-standard subjective practice but in a quantitative and intuitive manner.

Although computational annotation is convenient, manual annotation remains most accurate for many applications. Targeted, high-fidelity, manual annotation of specific features permits hypothesis-driven interrogation using spatial point pattern landscape analysis. A spatial point pattern of marked features in 2-dimensional space allows simple measures of feature density and distances to be calculated, and the clustering and dispersal of annotated features can be quantified by well-characterised specialised mathematical functions (Figure 2a)^9^.

**Figure 2.**
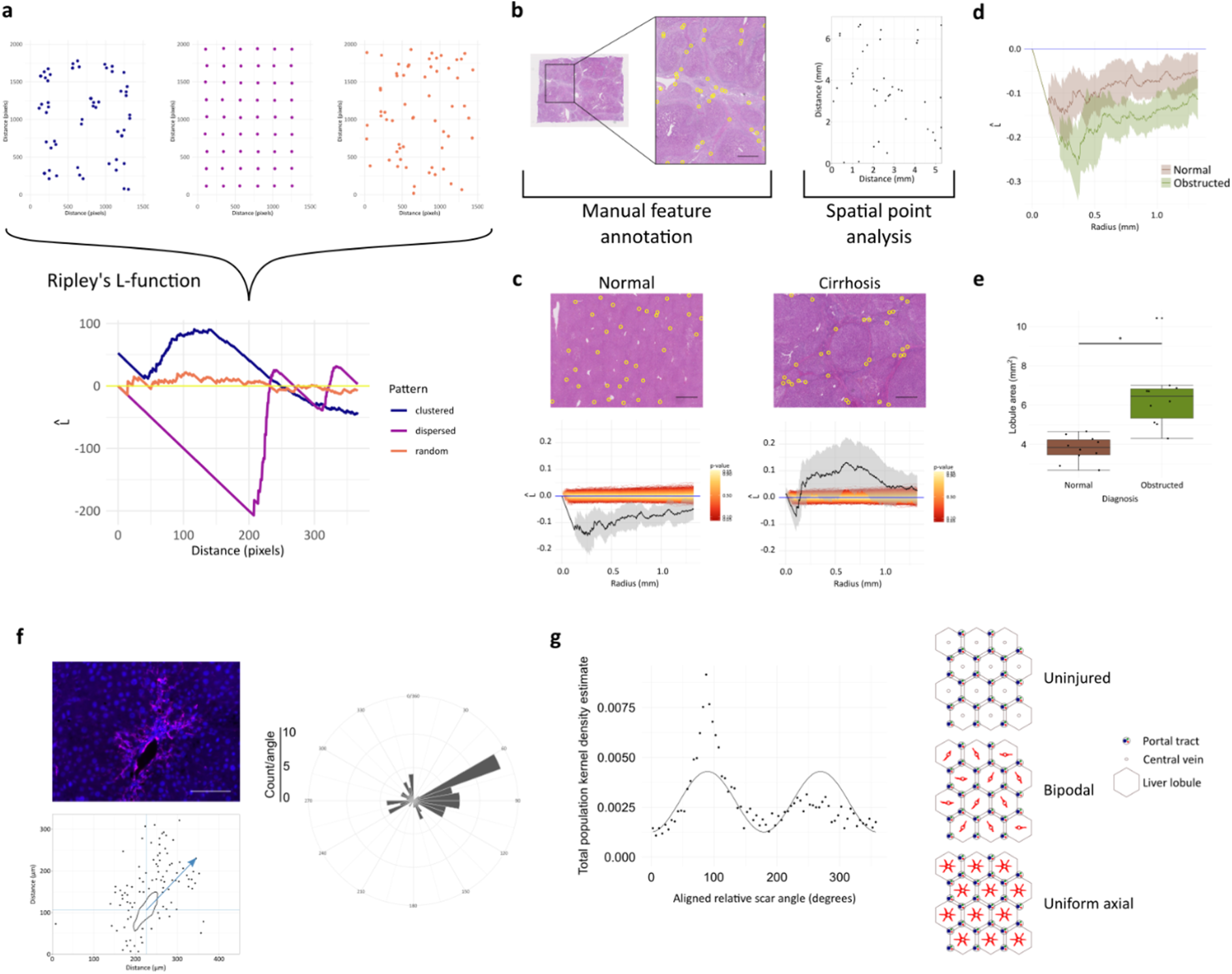
Spatial point pattern analysis of discrete features quantifies occult pathological feature relationships. A. Ripley’s L-function describes the organisation of features to quantify clustering or dispersal; clustered features are above, and dispersed features below, the horizontal line (yellow) representing complete spatial randomness. B. Example annotation of portal tracts by identification of hepatic artery branches in human liver, and plotted companion points. Scale bar 1 mm. C. Spatial point pattern analysis quantifies a traditional and previously subjective central tenet of liver disease, proving significant portal tract dispersal in normal liver that is lost in end-stage chronic liver disease (Ripley’s L-function with 95% confidence intervals, n=10). Scale bars 1 mm. D. In peripheral liver with subjectively normal architecture from in patients with hilar tumours, spatial point pattern analysis demonstrates that portal tracts are significantly more dispersed (Ripley’s L-function with 95% confidence intervals, n=10) with increased lobular size (E, data represented as individual points with median (centre line), first and third quartiles (lower and upper box limits), 1.5x interquartile range (whiskers), n=10, *Welch unpaired two-sample two-sided t-test p=0.0005835). F. Spatial point patterns of scar-orchestrating (α-smooth muscle actin positive) cells and central vein profiles after profibrotic injury allow the radial distributions of cells to be aligned to quantify the fundamental orientation of scar formation. Scale bar 100 μm. G. Liver scarring is organised with a dominant scarring axis accompanied by a single secondary directly-opposed axis rather than developing uniformly along all available central-central axes.

Spatial point patterns were derived from manual annotations of large vascular structures in images of normal liver and end-stage scarred (cirrhotic) liver and their distributions evaluated (Figure 2b). For liver pathologists, the loss of the regular hepatic architecture is the subjective histological *sine qua non* of end-stage liver disease. Landscape reinterpretation allowed this to be quantified and statistically validated for the first time (Figure 2c). A tendency towards clustering offers support for the development of cirrhosis through a mechanism of parenchymal extinction that ‘draws together’ adjacent structures^10^. The same approach was applied to peripheral liver from cases with central tumours, all clinically reported as having normal microarchitecture. The distribution of portal tracts in liver with central tumours demonstrated greater dispersal than in normal liver (Figure 2d). Additional annotation of central veins (Supplementary figure 2) allowed the size of liver lobules, a microarchitectural functional unit, to be modelled to demonstrate pathological but subjectively occult enlargement (Figure 2e). Thus, complementary landscape metrics readily defined previously unquantifiable and poorly recognised disease-related structural changes.

Image sets from obstructed and normal renal cortex (Supplementary figure 3), and normal pancreas (Supplementary figure 4), were also examined to demonstrate multi-organ applicability. Glomerular and Islet of Langerhans distributions, respectively, could be quantified by the same functions. The renal cortex in centrally obstructed kidneys did not demonstrate derangement of normal architecture equivalent to that found in the liver, suggesting fundamental differences in organ plasticity and response to injury.

The relational context of cells, as well as tertiary structures, can also be defined by a landscape approach to generate additional mechanistic insight. A dataset of images from a rodent model of early scarring in fatty liver disease was used for manual annotation of both the positions of scar-orchestrating myofibroblasts (MFBs)^11^ and the focal point of injury, central veins. Spatial point patterns of MFBs and the central vein circumference were used to define the relative cell positions. The distribution of MFB-to-central vein distances and relative scar axis, based on peak MFB density, was determined for each animal (Figure 2f). Examination of this fundamental disease process in relation to a fixed histological landmark revealed that scarring is initiated in a bipolar manner, rather than along all possible axes of scar development, indicating an unknown property of scar initiation (Figure 2g). The distribution of MFBs with respect to distance from the same landmark could also be calculated as a phenotypic feature (Supplementary figure 5).

Our framework depends upon a conceptual shift to consider a histological section as a tissue landscape, releasing the rich topography for interrogation by the methodologies of geosciences and landscape ecology. In this manner, features describing normality can be quantified to permit deviation from these norms to be identified. The approach can leverage histologically fluent manual annotation in hypothesis-driven work, limited only by creativity. The user-specified suite of metrics describes previously unquantifiable feature relationships over all microarchitectural scales. Critically, given the proliferation of computational methods to quantify images, this approach to a fully segmented image permits a complete suite of new metrics to be generated in a species-, tissue-, disease-, or segmentation methodology-agnostic manner without any additional training requirement.

## Figures

**Supplementary figure 1.**
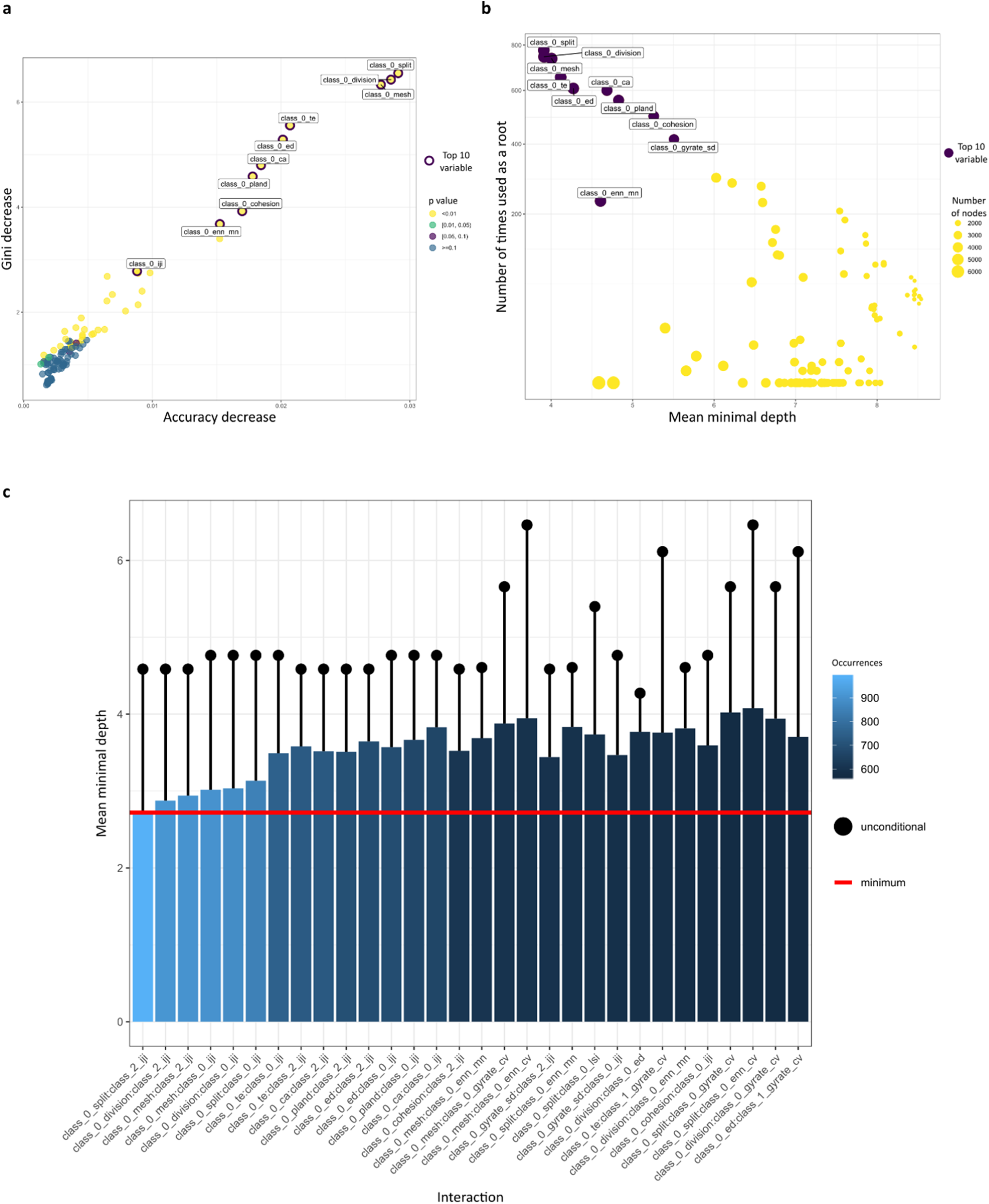
The suite of landscape metrics generated from fully segmented images can be used for machine-learning disease classification. Fully segmented images were generated from paired hepatocellular carcinoma or non-lesional liver using a simple WEKA haematoxylin and eosin stain deconvolution classifier identifying nuclei, cytoplasm and vasculature. Images were used in a patch landscape analytic pipeline, and the FRAGSTATS suite of metrics and other landscape complexity measures were generated. Metrics were used to train a random forest classifier to predict diagnosis. The individual top-ranked metrics by multiple measures; A. Accuracy decrease – mean decrease of prediction accuracy after variable is permuted, Gini decrease – mean decrease in the Gini index of node impurity by splits on variable, p-value – p-value of test determining whether the observed number of successes (number of nodes in which variable was used for splitting) exceeds the theoretical number of random successes. B. Mean minimal depth of variable, Number of times used as a root -total number of trees in which variable is used for splitting the root node, Number of nodes -total number of nodes that use variable for splitting. C. Interactions between nuclear features and those of the sinusoidal vasculature were most important/frequent (class 0 -nuclei, class 1 -cytoplasm, class 2 -vascular channels).

**Supplementary figure 2.**
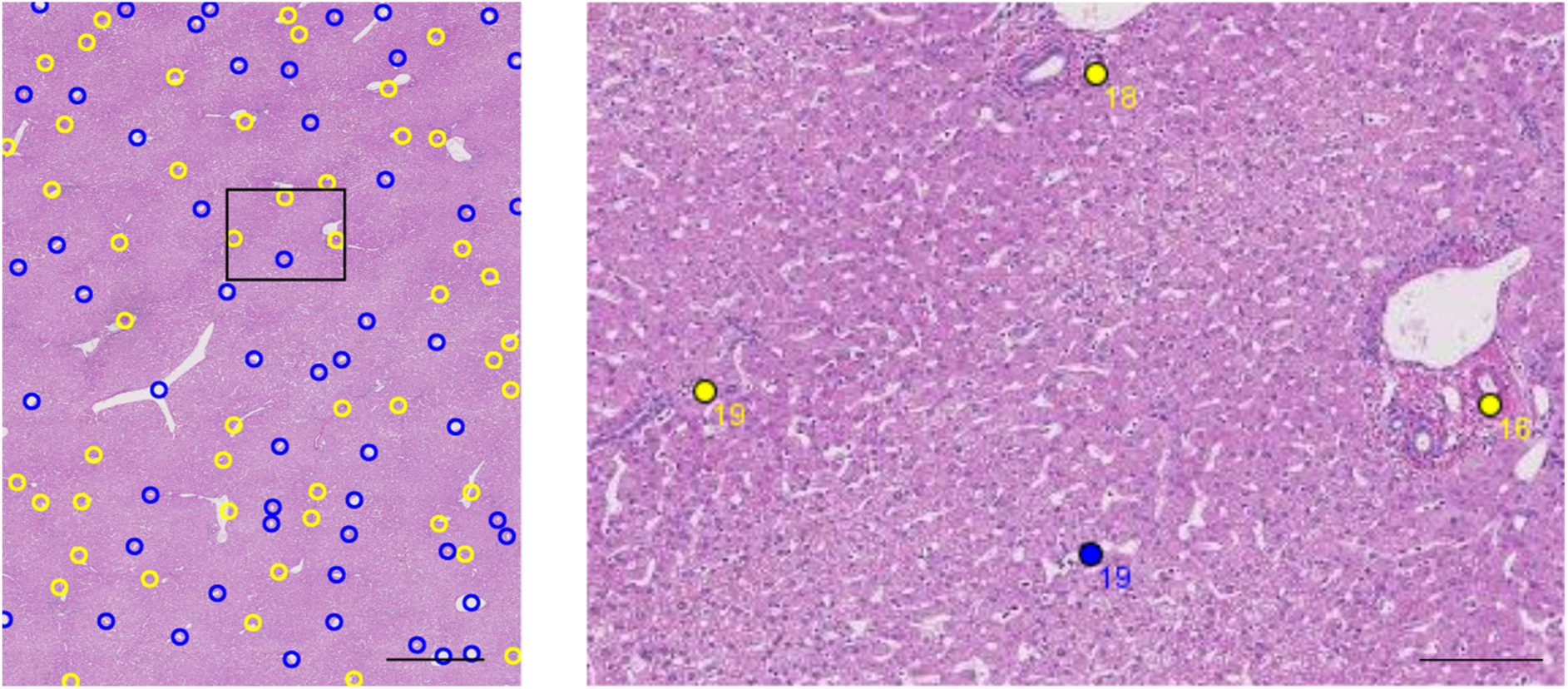
Portal-central vascular relationships can be quantified by separate annotations. In qualitatively normal liver, portal tract (yellow, identified by the presence of a hepatic artery branch) and central vein profiles (blue) can be separately annotated to allow the separate spatial point patterns to be interrogated to model lobule size and organisation. Scale bars 1 mm (left) and 100 μm (right).

**Supplementary figure 3.**
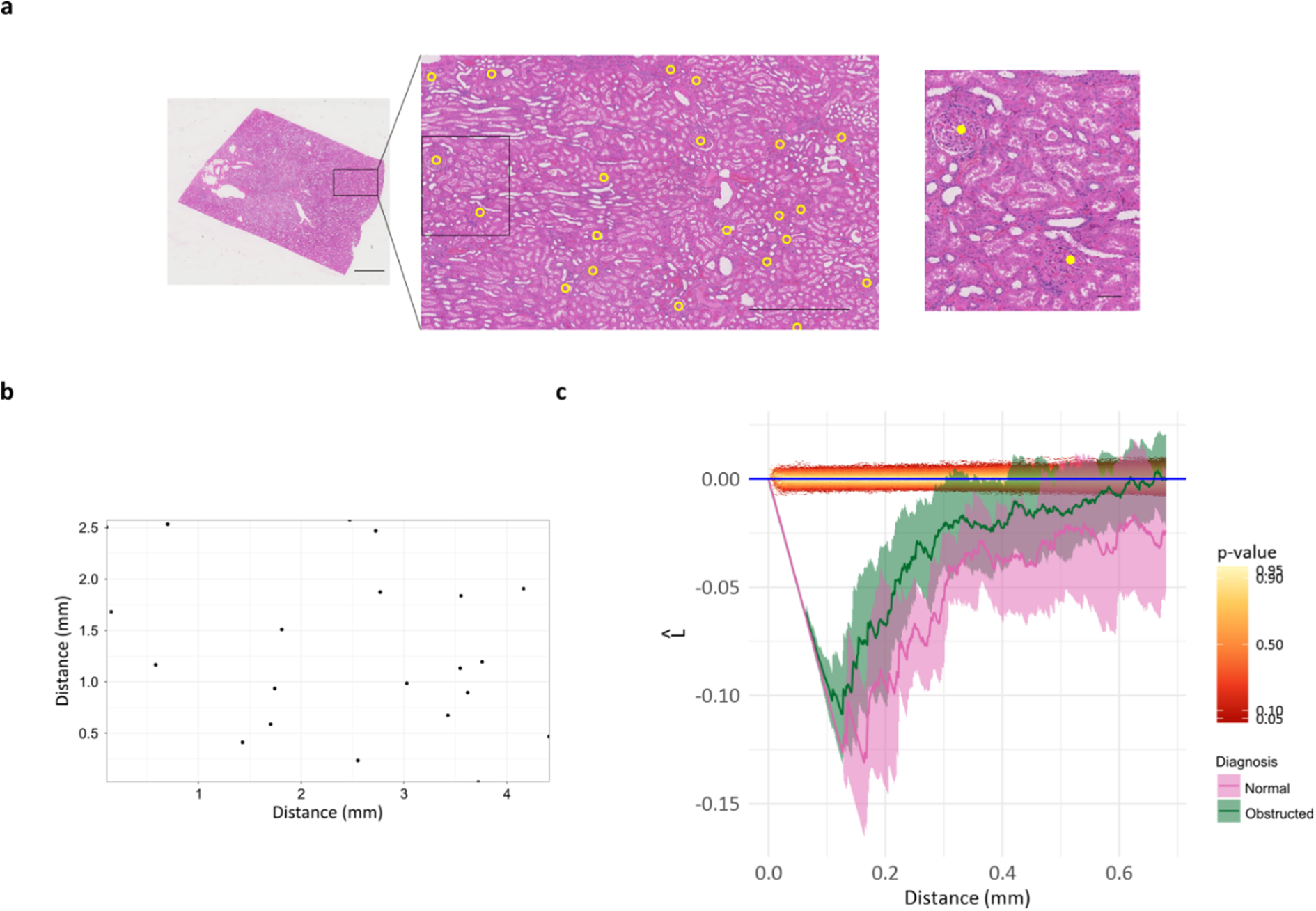
Renal cortical landscape assessment by spatial point pattern analysis. A. Fields from the cortex of normal kidney and from the kidneys with tumours of the renal pelvis were annotated to mark the glomeruli. Scale bars 3 mm left, 1 mm centre, 100 μm right. B. Glomerular positions were used to create spatial point patterns. C. Groupwise comparisons of Ripley’s L-function demonstrate glomerular dispersal that is unchanged in obstructed organs (Ripley’s L-function with 95% confidence intervals, n=8).

**Supplementary figure 4.**
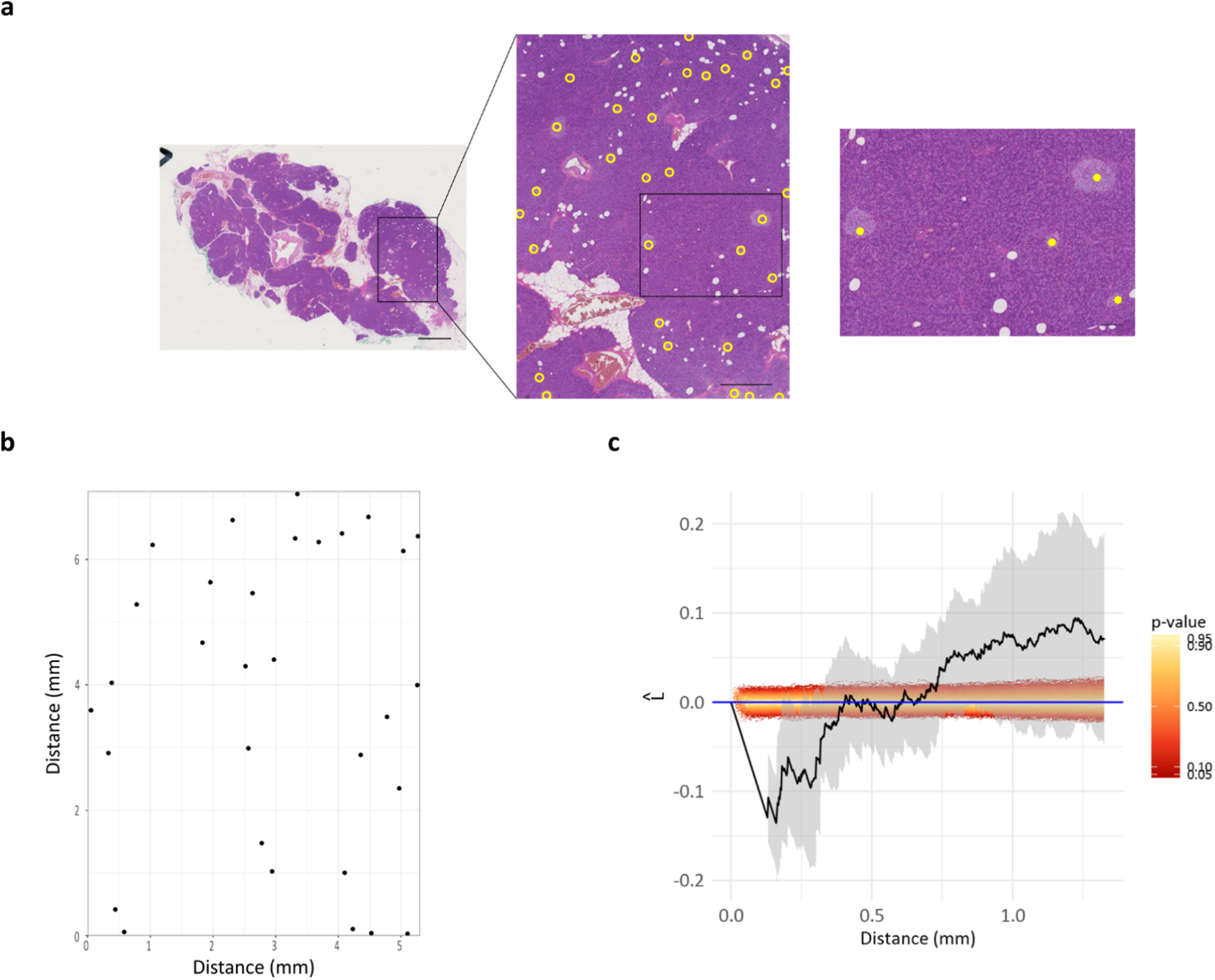
Pancreatic landscape assessment by spatial point patterns. A. Islets of Langerhans were annotated in images of normal pancreas and used to create spatial point patterns (B). Scale bars 3 mm left, 1 mm centre, 200 μm right C. The distribution of islets throughout the pancreas was not statistically separable from complete spatial randomness by Ripley’s L-function evaluation, n=10.

**Supplementary figure 5.**
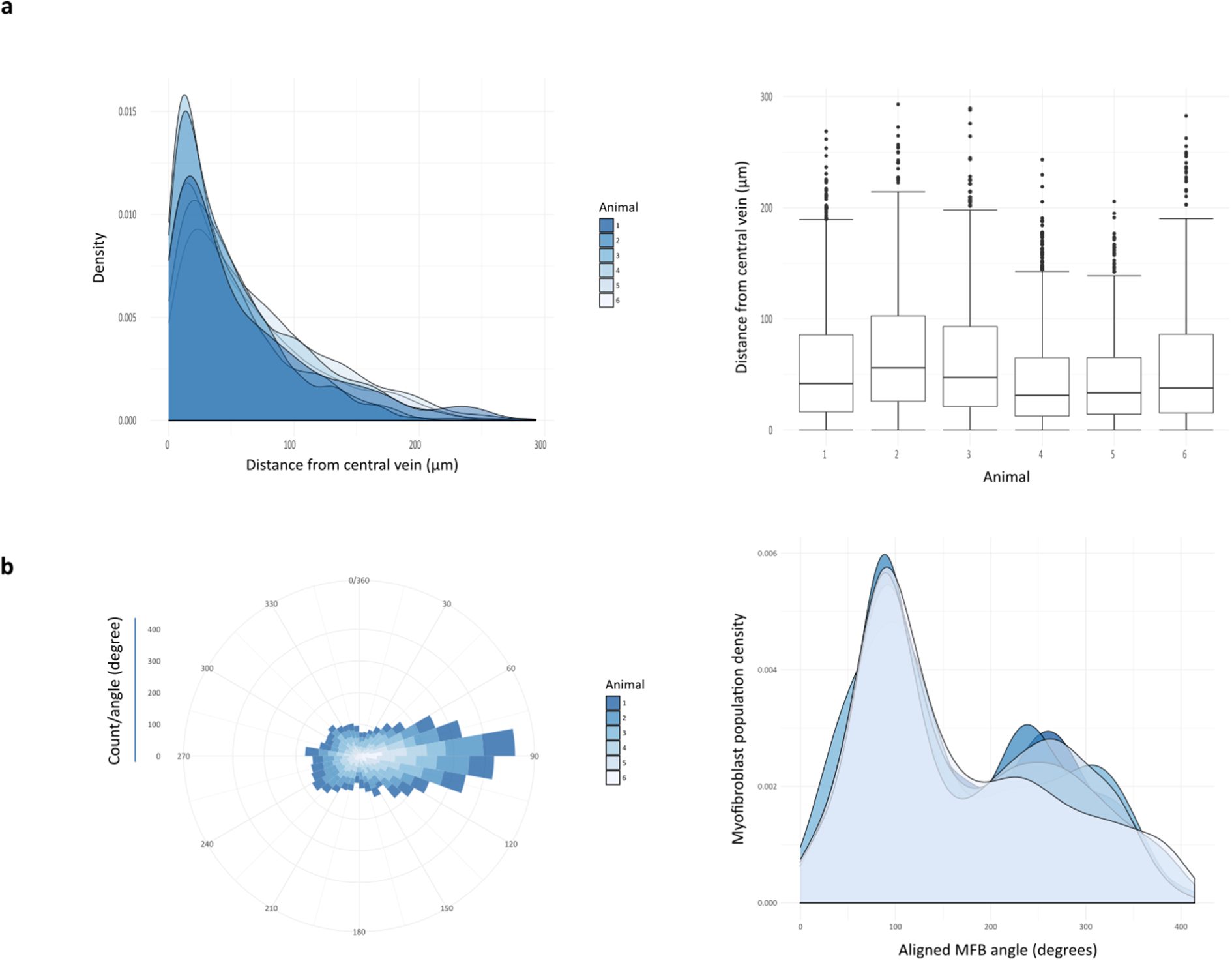
Landscape quantification of scar-orchestrating cell populations in disease models. A. The positions of scar-orchestrating cells in livers of mice chronically injured with CCl_4_ and the central vein profile were annotated, and separate spatial point patterns generated. The shortest distance of each cell from the central vein could be calculated, and the kernel density estimates (left) or individual distances for all cells (right, data represented with median (centre line), first and third quartiles (lower and upper box limits), 1.5x interquartile range (whiskers)) of each animal quantified. B. The angle of each cell with respect to the central vein centroid and an arbitrary pole could be calculated, and the axes aligned for each central vein profile based on the direction of the peak population density (left), and kernel density estimated of the aligned radial angle of each cell plotted for each animal.

## Methods

### Human tissue access

Human tissue was obtained by approved application to the Lothian NRS Human Annotated Bioresource that is authorised to provide unconsented anonymised tissue under ethical approval number 15/ES/0094 from the East of Scotland Research Ethics Service REC 1. All tissue was from cases from 2006 onwards and received anonymised to all details other than aetiology.

For manual annotation studies, single haematoxylin and eosin-stained sections from the deep hepatic parenchyma, sampled as part of the standard diagnostic specimen pathway, were used. No additional sections were required. Sections were obtained from cirrhotic explants with the 3 dominant patterns of fibrosis; primary biliary disease (n=11), steatohepatitis (n=10), and chronic Hepatitis C virus infection as a cause of lobular hepatitis (n=10). 10 non-lesional deep parenchymal blocks (> 5 cm from hilar lesional tissue) from liver with hilar cholangiocarcinoma were also obtained. 10 non-lesional liver sections from partial hepatectomies for metastatic disease (8 colorectal carcinomas, 1 melanoma) or a benign biliary cyst (single case) were used to represent normal liver.

8 cases of non-lesional pancreas from pancreaticoduodenectomies (Whipple’s procedure) for extrahepatic cholangiocarcinoma arising proximal to the confluence with the pancreatic duct were used to represent normal pancreas.

Routinely sampled blocks of non-lesional renal cortex from nephrectomies from 10 cases of conventional clear cell renal cell carcinoma, representing normal renal cortex and analogous to non-lesional blocks from partial hepatectomies for intrahepatic mass lesions, and from 10 cases ureteric or renal pyloric urothelial carcinoma, analogous to the hilar cholangiocarcinomas, were used.

For automated segmentation, single haematoxylin and eosin-stained sections including lesional (hepatocellular carcinoma) and adjacent non-lesional liver were obtained from 54 explants or resections containing hepatocellular carcinoma, without selection for aetiology or tumour grade.

### Murine model of liver fibrosis

Liver fibrosis was induced in cohorts of wild type C57Bl6 male mice by 8 weeks Carbon Tetrachloride (CCl_4_) injection twice weekly, 0.25 µl/g body weight in a 1:3 ratio with sterile olive oil^12^ or vehicle alone. Animals were not randomised to injury or control groups. Blinding to control or injury groups was not possible as injury is macroscopically and microscopically apparent. Animals were housed in a specific pathogen-free environment and kept under standard conditions with a 12 h day/night cycle and access to food and water ad libitum. All animal experiments were carried out under procedural guidelines, severity protocols and with ethical approval from the University of Edinburgh Animal Welfare and Ethical Review Body (AWERB) and the Home Office (UK).

### Scanning and image generation methods

Whole slide images of haematoxylin and eosin-stained human sections in .ndpi format were acquired using a Hamamatsu NanoZoomer to x20 depth. Tiled-TIFF thumbnails were generated from the .ndpi files using ndpisplit from the NDPITools suite^13^, and tiled-TIFF files converted to standard TIFF (for automated segmentation) or JPEG (for manual annotation) format compatible with ImageJ ^14^ by command-line ImageMagick.

### Immunofluorescence methods

Antigen retrieval of murine sections was achieved by microwaving in Tris-EDTA pH 9.0 for 15 minutes.

For immunofluorescent staining of aSMA, sections of murine liver were labelled with a monoclonal mouse antibody (Sigma A2547, clone 1A4, 1:1500 dilution, 1-hour incubation at room temperature). Staining was visualized with donkey anti-mouse IgG (H and L) Alexa Fluor 555 conjugated secondary antibody (ThermoFisher Scientific), and sections mounted in VECTASHIELD HardSet Antifade Mounting Medium with DAPI (Vector Laboratories). Negative controls were performed using identical concentrations of species and isotype-matched non-immune immunoglobulin in place of primary antibody or omission of primary antibody.

10 ×20 objective fields centred on a central vein (in keeping with the pattern of damage of CCl_4_) were acquired using a Zeiss Axioplan II microscope and Photometrics CoolSNAP HQ2 camera, and separate TIFF images of each channel exported.

### Manual identification and annotation of histological features

For human liver tissue, a central 5.32 mm × 7.11 mm (37.8 mm^2^) rectangular field from each .jpeg thumbnail whole slide image, the largest that could be taken from every scan, was cropped in FIJI^15^ and used to mark, as separate region of interest (ROI) sets, the centre of each central vein (from normal or centrally obstructed) and centre of each hepatic artery (identifying portal tracts when paired with a portal vein branch and/or bile duct). Marking was informed by viewing the WSIs in NDPIviewer (Hamamatsu) alongside to allow accurate identification.

For human pancreatic tissue, a central 5.32 mm × 7.11 mm (37.8 mm^2^) rectangular field from each .jpeg thumbnail whole slide image was used to mark the centre of each islet of Langerhans.

For human kidney, a 4.54 mm × 2.72 mm (12.35 mm^2^) rectangular field of renal cortex from each .jpeg thumbnail whole slide image was used to mark the centre of each glomerulus.

For murine myofibroblast (MFB) images, multichannel images were created in FIJI using the Image5D plugin, and the nucleus of each aSMA-positive MFB marked manually as a ROI set, excluding nuclei of concentrically arranged smooth muscle cells in vessel walls. The circumference of the central vein lumen also marked as a separate line segment ROI.

### Computational image segmentation

1mm^2^ ROIs from lesional (HCC) and non-lesional liver from each resection or explant case were selected manually and used to create 4 contiguous tiles from each.

A WEKA machine-learning classifier was trained in FIJI by a specialist liver transplant pathologist at the national liver transplant centre to simply deconvolve the staining into haematoxylin (nuclei), eosin (cytoplasm) and unstained areas (sinusoids/vessels). The classifier was applied to all tiles using a script that generated a classified TIFF output image.

### Spatial point pattern generation and analysis

Spatial point pattern and statistical analysis were undertaken in RStudio^16^. For each image, FIJI generated ROIs were imported using the *RImageJROI package*^17^ read.ijroi() function, and converted into *spatstat* package^18^ spatial point patterns using the ij2spatstat() function.

Spatial point pattern analysis was performed using the spatstat package. For distribution analysis of tertiary and quaternary structures in human tissue (portal tracts, central veins, islets of Langerhans, glomeruli), Ripley’s L-function^9^ was implemented with the Lest() function with the default edge corrections (Ripley’s isotropic, translation and border) applied; global envelopes using Monte-Carlo simulations of the theoretical L-function of complete spatial randomness (CSR) were derived by the envelope() function. Ripley’s L-functions of groups were compared with the studpermu.test() function.

To estimate individual lobule size based on the classical lobule depiction as a regular hexagon in normal and obstructed human liver, the distances from each central vein to the 6 nearest portal tracts were calculated with the nndist() function. For each central vein, the mean to the 6 distances (r) was used to calculate the area of the lobule 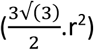.

For analysis of central vein-MFB radial distances, the nncross() function was used to determine the shortest distance to the central vein circumference for each aSMA-positive cell nucleus. For MFB directional analysis, the centroid of the central vein for each image was calculated using the centroid.owin() function, and used as (0,0). The position of each MFB was converted to polar coordinates to calculate the angle (ϕ_i_) from an arbitrary reference. Kernel density estimation of all MFB ϕ_i_ for each image was calculated with the density() function of the core stats package, and the angle of peak density (ϕ_peak_) determined. To allow comparison with distributions of MFBs from other images, all MFBs were effectively rotated about the central vein centroid such that ϕ_peak_ was 90°.

### Patch-based landscape analytics

Classified TIFF output images from WEKA/FIJI were used in a pipeline in RStudio that first converted each to a GeoTIFF image using the Universal Transverse Mercator projection and World Geodetic System (WGS) 84 datum using *rgdal* package accessing the Geospatial Data Abstraction Library^19^ and PROJ.4^20^. GeoTIFF images were used as input for the *landscapemetrics* package^21^ to analyse the categorical landscape patterns using metrics based on the FRAGSTATS suite^7^ as well as more recently developed measures of landscape complexity^8^.

### Machine learning disease classification

The paired HCC and non-lesional classified image set was used. Eighty per cent of cases were randomly chosen as a training set and the remainder used only as a validation set.

Landscape and class level metrics of the ‘aggregation’, ‘area and edge’, ‘diversity’, and ‘complexity’ groups were used as features for model training after near-zero variance features were removed using *caret::nearZeroVar*^22^. Features of the training set were optimally normalised using *bestNormalize*^23^, and features selected for model training by removal of those that were highly correlated (>0.75). A random forest model with 10000 trees was constructed to predict disease classification (HCC or non-lesional) using *randomForest*^24^. Variable importance measures of the constructed forest^25,26^ were calculated using *randomForestExplainer.*

### Third-party images

A satellite image from the European Space Agency Copernicus Sentinel-2B satellite L1C 2019-02-26 dataset was retrieved using the Sentinel Hub EO Browser under CC BY 4.0. The corresponding mapped region was retrieved from OpenStreetMap and used under CC BY-SA 2.0 (© OpenStreetMap contributors) to generate the composite image.

### Statistical methods

Distributions of MFB subpopulations were evaluated with a bootstrap version of the Kolmogorov– Smirnov test, ks.boot(), in the Matching package^27^.

For inter-group comparison of lobular area and central vein-MFB distances, normality of data was determined using Shapiro-Wilk testing and by examination of qq plots. After assumptions of normality were satisfied, the Welch (unequal variance) t-test was used to compare two groups^28^.

## Data availability statement

The datasets generated during and/or analysed during the current study are available from the corresponding author on reasonable request.

## Code availability statement

R scripts used to implement packages documented/cited are available from the corresponding author upon reasonable request.

## Acknowledgements

T.J.K. received a Wellcome Trust Intermediate Clinical Fellowship (095898/Z/11/Z).

## Author contributions

Conceptualization, T.J.K., A.M.T., J.P.I.; Methodology, T.J.K.; Formal Analysis, T.J.K.; Investigation, T.J.K., C.D.; Writing – Original Draft, T.J.K.; Writing – Review & Editing, T.J.K., C.D., A.M.T., J.P.I.; Funding Acquisition, T.J.K., J.P.I.

